# Spectral CytoFRET2 identifies lysine acetyltransferase inhibitors as modulators of vimentin assembly

**DOI:** 10.64898/2026.07.08.737169

**Authors:** Frédéric Larbret, Marie Irondelle, Sophie Tartare-Deckert, Marcel Deckert

## Abstract

Cytoskeletal plasticity is a defining feature of cancer progression, enabling tumor cells to adapt their morphology, mechanics, and migratory behavior during invasion and metastasis. Although actin filaments, microtubules, and intermediate filaments are known to cooperate in these processes, the molecular mechanisms coordinating their dynamics remain incompletely understood, particularly the role of post-translational modifications (PTMs). Here, we developed CytoFRET2, a multiparametric cytometry-based FRET platform that enables real-time and simultaneous monitoring of the dynamics of actin filaments, microtubules, and vimentin in living suspension cells. The system combines fluorescently tagged cytoskeletal reporters with spectral flow cytometry, allowing simultaneous high-content analysis of multiple cytoskeletal networks while overcoming autofluorescence and fluorescence interference from small molecules. Using spectral CytoFRET2, we screened a small library of epigenetic compounds targeting regulators of acetylation and methylation pathways. The screen revealed that inhibition of lysine deacetylases (KDACs) and sirtuins promoted stabilization of both microtubules and vimentin filaments, without impacting actin filament organization. In contrast, inhibition of lysine acetyltransferases (KATs), particularly with garcinol and anacardic acid, induced rapid vimentin disassembly. Mechanistically, the study reveals acetylation as a key post-translational modification regulating the dynamics of microtubules and vimentin filaments, with KAT inhibitors emerging as potent modulators of vimentin organization. Together, the findings establish spectral CytoFRET2 as a versatile platform for systematic investigation of cytoskeletal regulatory networks and therapeutic vulnerabilities in cancer.

## INTRODUCTION

Cytoskeletal plasticity is a hallmark of malignant transformation, enabling tumor cells to adapt their morphology, mechanics, and migratory behavior to dynamic microenvironmental constraints ^1,2^. Through coordinated remodeling of actin filaments, microtubules, and intermediate filaments, cancer cells acquire invasive capabilities, withstand mechanical stress, and disseminate to distant organs ^3^. Although each cytoskeletal network has been extensively studied individually, increasing evidence indicates that their functional integration is essential for tumor progression and metastasis ^4,5^. However, the molecular mechanisms coordinating these interconnected filament systems remain poorly understood.

A major layer of cytoskeletal regulation is mediated by post-translational modifications (PTMs) of cytoskeletal proteins and their associated regulators. Acetylation, phosphorylation, and other PTMs influence filament assembly, stability, and intermolecular interactions, thereby shaping cytoskeletal architecture and dynamics ^6,7^. In particular, α-tubulin acetylation has emerged as a critical determinant of microtubule stability, intracellular trafficking, and mechanotransduction ^6,8^. In cancer, widespread dysregulation of PTM-regulating enzymes, including lysine acetyltransferases (KATs), lysine deacetylases (KDACs), and sirtuins, suggests that aberrant PTM signaling may directly reprogram cytoskeletal organization and cellular mechanics ^9,10,11^. Nevertheless, establishing causal links between specific PTMs and coordinated remodeling of distinct cytoskeletal networks has remained challenging.

Among intermediate filaments, vimentin is a key regulator of cellular plasticity during epithelial–mesenchymal transition, invasion, and metastatic dissemination ^12,13^. Vimentin organization and dynamics are tightly controlled by phosphorylation and other PTMs ^14,15^, and studies have highlighted functional crosstalk between vimentin filaments and microtubules ^16,17^. However, how PTM-dependent microtubule dynamics influence vimentin assembly and subcellular organization remains largely unresolved, in part due to the lack of experimental platforms capable of dynamically studying multiple cytoskeletal networks on a large scale.

Current imaging-based approaches provide high spatial resolution but are inherently low throughput and difficult to adapt to systematic perturbation screens. Conversely, flow cytometry enables quantitative population-level analysis but has remained underexploited for studying cytoskeletal filament dynamics because of technical limitations related to fluorescent compounds, spectral overlap, and cellular autofluorescence ^18,19^. These limitations have hindered the unbiased identification of PTM-dependent regulators of cytoskeletal plasticity and restricted exploration of their therapeutic relevance.

Here, we address these challenges by developing CytoFRET2, a cytometry-based FRET platform that enables real-time, multiparametric analysis of actin, microtubule, and vimentin dynamics in living suspension cells. By integrating CytoFRET2 with spectral flow cytometry ^20^, we overcome fluorescence interference and autofluorescence-related artifacts, enabling high-throughput screening of PTM-targeted compound libraries.

Using this approach, we identified pan-KDAC inhibitors and KAT inhibitors as key regulators of microtubule and vimentin filament organization, revealing acetylation as a central protein modification controlling the assembly and disassembly of these filaments. Together, our findings uncover a novel regulatory axis governing cytoskeletal plasticity and establish spectral CytoFRET2 as a potent tool for systematic dissection of cytoskeletal regulatory networks with therapeutic potential.

## MATERIALS AND METHODS

### Cells and culture conditions

The human T leukemic cell line Jurkat (clone E6.1; ATCC TIB-152) was cultured in RPMI-1640 medium supplemented with 10% fetal bovine serum (FBS; HyClone). HEK293T cells (ATCC CRL-3216) were cultured in DMEM 10% FBS. Cells were maintained at 37 °C in a humidified incubator containing 5% CO₂.

### Chemical library and inhibitors

To modulate protein post-translational modifications, we used the SCREEN-WELL® Epigenetics Library (Enzo Life Sciences; BML-2836-0100), containing 41 small molecules targeting major epigenetic regulators. The library includes inhibitors of histone deacetylases (KDACs), sirtuins (SIRTs), lysine demethylases, histone acetyltransferases (KATs), histone methyltransferases, and DNA methyltransferases, as well as activators of sirtuins. Other inhibitors used in this study were from Merck.

### Plasmid construction

Plasmid design, cloning, and sequence verification were performed using standard molecular biology procedures. The mOrange plasmid was obtained from Clontech, CFP-H148D was provided by Dr. O. Albagli ^21^, and the T-Sapphire construct was provided by Dr. Zapata-Hommer. The ACT reporter cell line was generated as previously described ^22^. The TUB, VIM, and CT reporter lines were generated by lentiviral transduction using vectors derived from pRRLSIN.cPPT.PGK-EGFP.WPRE (Addgene #12252). The EGFP coding sequence was replaced with cDNAs encoding mOrange, CFP-H148D, or T-Sapphire using BamHI and BsrGI restriction sites. For α-tubulin fusion constructs, α-tubulin cDNA (Clontech #6324488) was amplified by PCR using the following primers: forward, 5ʹ-ATGAGCTGTACAAGTCCGGACT-3ʹ; reverse, 5ʹ-GTCGACTTAGTATTCCTCTCCTTCTTC-3ʹ. The PCR product was digested with BsrGI and SalI and inserted into the modified lentiviral vectors. For vimentin fusion constructs, vimentin cDNA was excised from pVimentin-PSmOrange (Addgene #31922) using AgeI and BglII and subcloned into pRRLSIN.cPPT.PGK-EGFP.WPRE and pRRLSIN.cPPT.PGK-mOrange.WPRE using AgeI and BamHI restriction sites.

### Lentivirus production

Lentiviral particles were produced as previously described ^23^. Briefly, HEK293T cells were co-transfected with the packaging plasmids pMD2.G (Addgene #12259) and pCMVR8.74 (Addgene #22036), together with the corresponding pRRL-derived transfer vector, using Lipofectamine 2000 (Invitrogen). Seventy-two hours after transfection, supernatants were collected, centrifuged at 3,000 rpm for 5 min to remove cellular debris, and filtered. Viral suspensions, estimated to contain approximately 10⁹ infectious units/mL, were aliquoted and stored at −80 °C until use.

### Generation of FRET reporter cell lines

To generate the FRET reporter cell lines, 1 × 10⁵ Jurkat cells were incubated for 12 h with 1 mL of lentiviral supernatant encoding either CFP-H148D or T-Sapphire. Cells were subsequently washed and cultured for an additional 48 h before single-cell sorting using a BD FACSAria II cell sorter. After approximately 3 weeks of expansion, CFP-H148D–expressing clones were transduced with EGFP-actin and mOrange-actin constructs, whereas T-Sapphire–expressing clones were transduced with Vimentin-EGFP and Vimentin-mOrange vectors. Wild-type Jurkat cells were independently transduced with EGFP-tubulin and mOrange-tubulin constructs. A control reporter line was generated by co-expressing unfused EGFP, mOrange, CFP-H148D, and T-Sapphire fluorescent proteins. Cells exhibiting balanced and robust EGFP and mOrange expression were isolated by cell sorting to establish four stable reporter cell lines: ACT (actin), VIM (vimentin), TUB (microtubules), and CT (control).

### Screening protocol

The four CytoFRET2 Jurkat reporter cell lines were co-seeded in 48-well plates at a density of 1 × 10⁵ cells/mL 24 h before analysis. Cells were treated for 1 h at 37 °C with individual compounds from the epigenetic library or DMSO as a control. For conventional flow cytometry, data were acquired using a FACSCanto II flow cytometer (BD Biosciences) equipped with 405-, 488-, and 633-nm lasers and an automated carousel loader. Each compound was tested in triplicate at 15, 50, and 100 µM. Twenty DMSO-treated wells were included in each experiment as internal controls. FRET efficiency was calculated using the following equation:

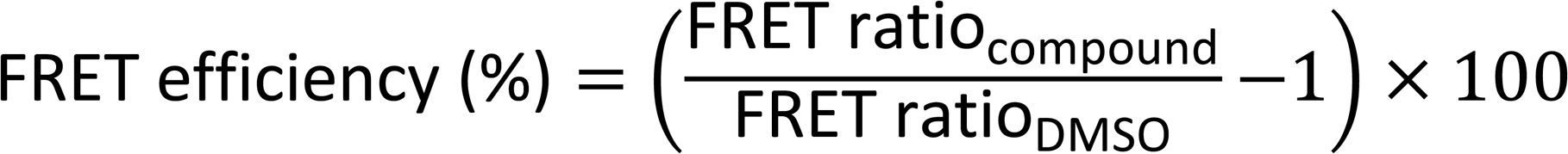

where the DMSO FRET ratio corresponds to the mean fluorescence intensity ratio obtained from all DMSO control wells within the same experiment.

### Flow cytometry FRET measurements

For conventional flow cytometry, cells were excited using the 488-nm laser. EGFP donor fluorescence was collected through a 530/30-nm bandpass filter, whereas mOrange FRET emission was detected using a 610/15-nm bandpass filter. The FRET ratio was calculated as the ratio of mOrange to EGFP fluorescence intensity following 488-nm excitation. CFP-H148D and T-Sapphire barcode signals were detected following 405-nm excitation using 485/45-nm and 530/30-nm bandpass filters, respectively, separated by a 502-nm long-pass dichroic mirror. Four barcode populations corresponding to ACT, VIM, TUB, and CT cells were identified based on CFP-H148D and T-Sapphire fluorescence profiles. At least 5,000 cells within the live single-cell gate were analyzed for each sample. For spectral flow cytometry, analyses were performed using an Aurora spectral cytometer (Cytek Biosciences) equipped with 355-, 405-, 488-, and 630-nm lasers. Spectral unmixing was performed using SpectroFlo software (Cytek Biosciences). Data from both conventional and spectral cytometry experiments were analyzed using FlowJo software (BD Biosciences).

### Confocal microscopy

Live cells were seeded onto chambered coverslips (Lab-Tek; Thermo Fisher Scientific) and imaged using a Zeiss Axiovert 200 M inverted fluorescence microscope equipped with a ×63 oil-immersion objective and a Hamamatsu Orca CCD camera. Acquisition settings were kept identical across all experimental conditions. When required, image stacks were deconvolved using Volocity software (Improvision).

### Statistical analysis

Unless otherwise indicated, experiments were independently repeated at least three times, and representative data or images are shown. Statistical analyses were performed using GraphPad Prism software (GraphPad Software). Data are presented as mean ± SD. Statistical significance was determined using Student’s *t*-test, with *P* < 0.05 considered statistically significant.

## RESULTS

### Validation of the CytoFRET2 platform

To extend the CytoFRET approach previously developed for monitoring actin dynamics ^22^, we generated CytoFRET2, a multiplexed FRET-based system enabling simultaneous analysis of actin filaments, microtubules, and vimentin intermediate filaments in living suspension cells (Figure 1A). CytoFRET2 consists of four Jurkat-derived reporter cell lines: ACT (actin), TUB (microtubules), VIM (vimentin), and a control line (CT). In the ACT, TUB, and VIM lines, EGFP and mOrange were fused to actin, α-tubulin, or vimentin, respectively, whereas the CT line co-expressed unfused fluorescent proteins (Supplementary Figure 1A). In this system, FRET efficiency reflects filament assembly state. Polymerization of fluorescently tagged cytoskeletal monomers increases the proximity between EGFP and mOrange, resulting in elevated FRET, whereas filament disassembly reduces the FRET signal. To enable multiplexed analysis, each reporter line was identified using a dual fluorescent barcode based on CFP and T-Sapphire expression: ACT (CFP⁺), VIM (T-Sapphire⁺), TUB (double negative), and CT (double positive). Following gating on live single cells, FRET ratios were quantified independently for each population. To validate the platform, CytoFRET2 cells were exposed to a panel of reference compounds targeting distinct cytoskeletal networks (Figure 1B). The microtubule stabilizer paclitaxel ^24^ increased FRET efficiency in TUB cells by 15–25%, whereas colchicine and related depolymerizing agents reduced FRET by 5–9%. Similarly, the actin stabilizers jasplakinolide ^25^ and WP1130 ^26^ increased FRET in ACT cells by up to 33%, while the actin depolymerizing agent latrunculin B ^27^ decreased FRET across the ACT, TUB, and VIM lines. Inhibition of PP2A using okadaic acid or calyculin A, both known to induce vimentin hyperphosphorylation and filament disassembly ^28,29^, selectively reduced FRET in VIM cells by 20–23%. In contrast, microtubule depolymerization induced a perinuclear redistribution of vimentin associated with increased FRET. Dose–response analyses (Figure 1C and Supplementary Figure 1B) and confocal microscopy (Figure 1D) confirmed these observations. Stabilizing compounds such as paclitaxel and jasplakinolide enhanced microtubule and actin filament organization respectively, whereas PP2A inhibition induced pronounced vimentin disassembly. Collectively, these results demonstrate that CytoFRET2 quantitatively reports the assembly state of actin, microtubule, and vimentin networks in living cells.

**Figure 1.**
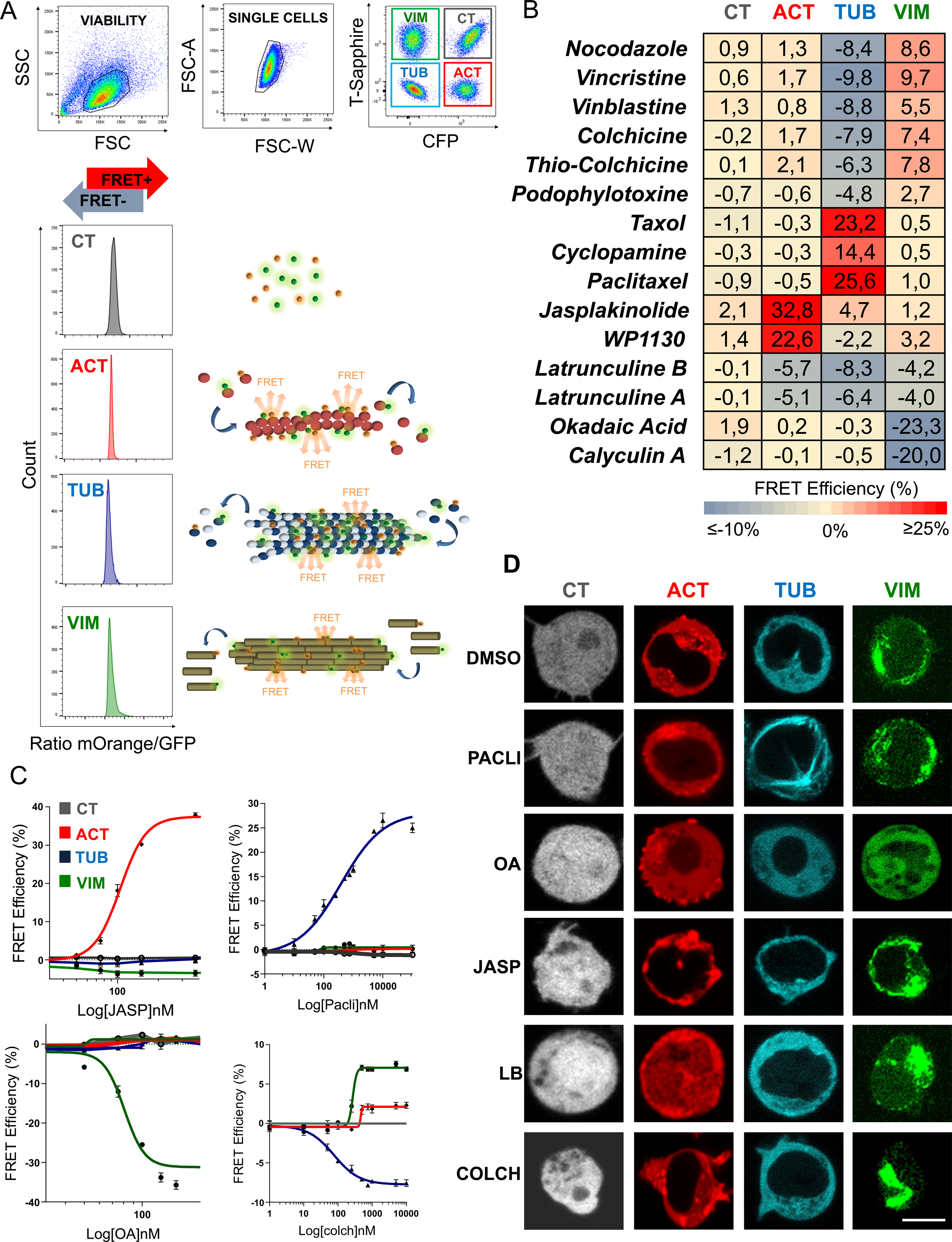
Principle and validation of the CytoFRET2 platform for multiplexed analysis of cytoskeletal dynamics. **(A)** Schematic overview of the CytoFRET2 assay. The platform combines four Jurkat-derived reporter cell lines: ACT (actin), TUB (microtubules), VIM (vimentin), and CT (control). In ACT, TUB, and VIM cells, GFP-and mOrange-tagged cytoskeletal proteins were expressed to monitor filament assembly by FRET, whereas CT cells co-expressed unfused fluorescent proteins as a negative control. Reporter populations were distinguished using a fluorescent barcoding strategy based on CFP and T-Sapphire expression : ACT cells (CFP⁺), VIM cells (T-Sapphire⁺), CT cells (CFP⁺/T-Sapphire⁺), and TUB cells (double negative). Representative flow cytometry plots illustrate the sequential gating strategy, including selection of live cells based on FSC/SSC parameters, singlet discrimination, and identification of each reporter population according to CFP and T-Sapphire fluorescence. Representative histograms show the mOrange/GFP fluorescence ratio used to quantify FRET efficiency. A rightward shift indicates increased FRET and filament assembly, whereas a leftward shift indicates reduced FRET and filament disassembly. **(B)** Heatmap summarizing CytoFRET2 analysis following treatment with a reference panel of cytoskeleton-targeting compounds. Cells were treated for 60 min before flow cytometry analysis. FRET efficiencies are expressed as percentage variation relative to DMSO-treated controls. **(C)** Dose–response analysis of FRET efficiency in ACT, TUB, and VIM reporter lines following treatment with jasplakinolide, paclitaxel, okadaic acid, or colchicine. Data represent mean ± SD from three independent experiments (*n* = 3). **(D)** Representative fluorescence microscopy images showing cytoskeletal organization in each reporter cell line following treatment with the indicated compounds.

### Epigenetic library screening identifies PTM-dependent regulators of cytoskeletal organization

To investigate how post-translational modifications regulate cytoskeletal dynamics, we combined CytoFRET2 with a pharmacological library targeting enzymes involved in acetylation and methylation pathways. The library included inhibitors of KDACs, KATs, and sirtuins, as well as modulators of DNMTs, HDMTs, LDMTs, and HMTs (Figure 2A and Supplementary Table 1). The four CytoFRET2 reporter lines were co-seeded and treated for 1 h with each compound at 15, 50, or 100 µM. FRET efficiencies were quantified by flow cytometry and expressed as percentage variation relative to DMSO-treated controls (Figure 2B). Significant changes were defined using the variability observed across 20 independent DMSO control wells (Supplementary Figure 2A). Among the 41 compounds tested, seven altered the FRET signal in the CT control line, indicating direct interference with fluorescent reporter signals. These compounds, mainly targeting sirtuins, were excluded from subsequent analyses. Of the remaining 34 compounds, 13 increased FRET efficiency in TUB cells, including eight HDAC inhibitors. Notably, five of these compounds also increased FRET in VIM cells, suggesting coordinated stabilization of microtubules and vimentin filaments. In contrast, two KAT inhibitors, garcinol and anacardic acid, markedly reduced FRET in VIM cells by 30–40% at 100 µM, consistent with vimentin filament disassembly. Garcinol induced this effect within 1 h, whereas anacardic acid required prolonged incubation (4 h) to produce detectable changes (Supplementary Figure 2B). Dose–response analyses further highlighted the strong activity of four HDAC inhibitors, Oxamflatin, CI994, MC1293, and TSA, all of which induced pronounced increases in FRET efficiency in both TUB and VIM cells (Figure 2C). Conversely, garcinol selectively reduced FRET in the VIM cell line. Together, these findings identify acetylation-dependent regulation as a major determinant of microtubule and vimentin organization, with KDAC inhibition promoting filament stabilization and KAT inhibition favoring vimentin disassembly.

**Figure 2.**
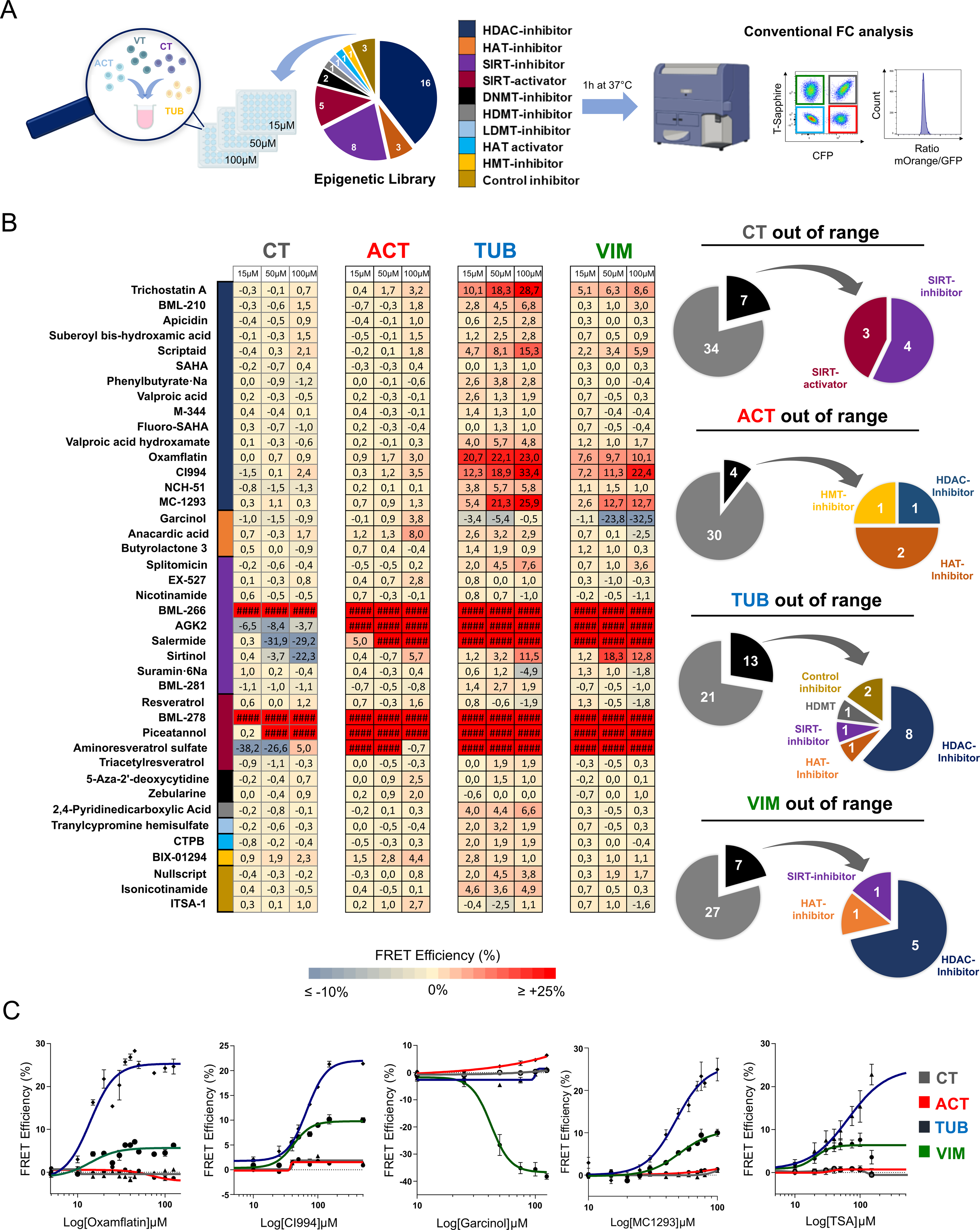
CytoFRET2 screening identifies acetylation-dependent regulators of cytoskeletal organization. **(A)** Experimental workflow for epigenetic compound library screening using CytoFRET2. The four reporter cell lines were mixed and seeded in 96-well plates before treatment for 60 min with compounds targeting major epigenetic regulators, including HDACs, HATs, sirtuins, DNMTs, HDMTs, LDMTs, and HMTs. FRET signals were quantified by conventional flow cytometry and normalized to DMSO-treated controls. **(B)** Heatmap representation of FRET efficiencies measured in ACT, TUB, VIM, and CT reporter lines following treatment with each compound at 15, 50, or 100 µM. Data are expressed as percentage variation relative to DMSO-treated controls and represent mean ± SD from three independent screens (*n* = 3). Grey and black ring charts indicate the number of compounds inducing FRET variations exceeding the threshold defined from DMSO controls. Colored ring charts classify compounds according to their epigenetic target family. **(C)** Dose–response curves showing FRET efficiencies in control (grey), ACT (red), TUB (blue), and VIM (green) reporter lines following treatment with Oxamflatin, CI-994, garcinol, MC1293, or trichostatin A (TSA).

### Spectral flow cytometry improves CytoFRET2 performance and resolves fluorescence interference

To overcome fluorescence interference caused by intrinsically fluorescent compounds, we adapted CytoFRET2 to spectral flow cytometry. Unlike conventional cytometry, which detects fluorescence within predefined wavelength windows, spectral cytometry records the full emission spectrum (400–829 nm), enabling improved fluorochrome discrimination and separation of autofluorescent signals ^30^. We first validated the spectral workflow using non-fluorescent reference compounds previously analyzed by conventional cytometry. CytoFRET2 analysis by spectral cytometry included FSC/SSC gating and barcode discrimination based on CFP and T-Sapphire fluorescence (Figure 3A). Two unmixing strategies were compared, with or without autofluorescence extraction. Removal of autofluorescence substantially improved barcode resolution and separation of the four reporter populations. Spectral profiles generated using SpectroFlo revealed distinct emission signatures for each fluorescent protein and for cellular autofluorescence across all laser configurations. Importantly, FRET efficiencies obtained by spectral cytometry closely matched those measured by conventional cytometry across all reference compounds and reporter lines (Figure 3B), confirming the robustness of the approach. To further validate spectral FRET detection, fluorescence emission profiles were analyzed in ACT and TUB cells treated with DMSO or cytoskeletal modulators (Figures 3C–D). Stabilizing compounds, including paclitaxel and jasplakinolide, increased mOrange emission (Aurora spectral detectors B5–B14; 588–829 nm) while reducing EGFP emission (Aurora spectral detectors B1–B4; 498–590 nm), consistent with enhanced FRET. Conversely, destabilizing agents such as colchicine and latrunculin B induced the opposite fluorescence shift, reflecting reduced FRET efficiency. Similar spectral changes were observed in VIM cells following okadaic acid treatment (Supplementary Figure 3). These results demonstrate that spectral flow cytometry provides FRET quantification and barcode discrimination comparable to conventional cytometry while substantially improving signal resolution through autofluorescence extraction.

**Figure 3.**
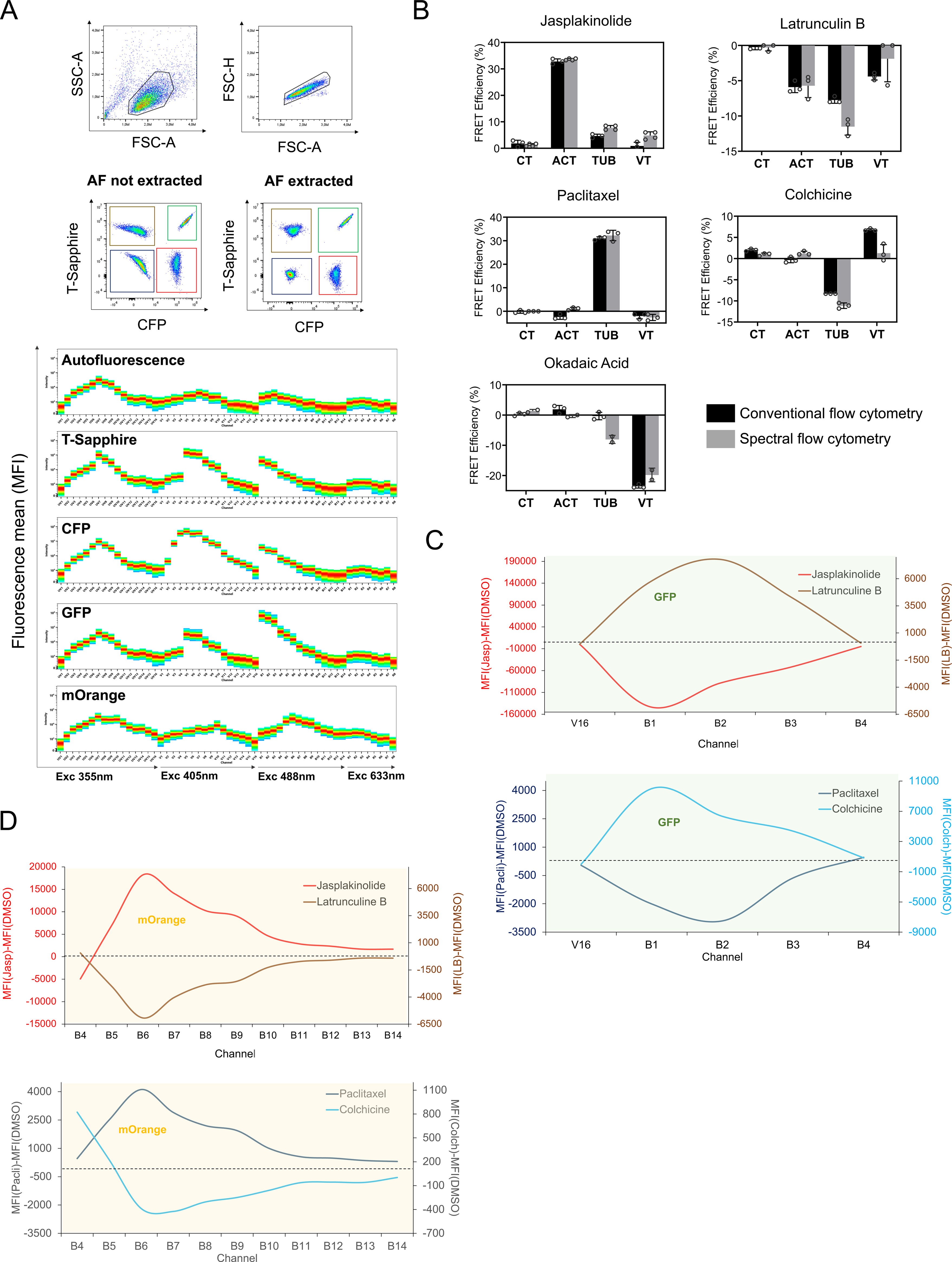
Validation of CytoFRET2 using spectral flow cytometry. **(A)** Spectral flow cytometry workflow for CytoFRET2 analysis. Representative dot plots illustrate gating of live single cells and fluorescent barcoding–based discrimination of ACT, TUB, VIM, and CT reporter populations in both conventional and spectral flow cytometry. Emission spectra of CFP, T-Sapphire, GFP, and mOrange are shown. Spectral analyses were performed using an Aurora spectral flow cytometer (Cytek Biosciences) equipped with 355-, 405-, 488-, and 633-nm lasers. **(B)** Comparison of FRET efficiencies measured by conventional and spectral flow cytometry following treatment with jasplakinolide (150 nM), paclitaxel (10 µM), okadaic acid (150 nM), or colchicine (1 µM) for 1 h at 37 °C. Black bars represent conventional flow cytometry measurements and grey bars represent spectral flow cytometry measurements. **(C)** Representative GFP emission spectra associated with FRET modulation induced by jasplakinolide, latrunculin B, paclitaxel, or colchicine treatment. **(D)** Representative mOrange emission spectra corresponding to the treatments shown in panel C.

### Spectral CytoFRET2 enables analysis of fluorescent compounds

We next re-evaluated the seven compounds excluded from the initial screen because of fluorescence interference. For each compound, the intrinsic fluorescence spectrum was incorporated into the spectral unmixing matrix as an independent fluorochrome (Supplementary Figure 4A). Spectral overlap analysis yielded complexity indices below 6 for all compounds (Supplementary Figure 4B), consistent with Cytek deconvolution guidelines. Spectral cytometry markedly improved CFP/T-Sapphire discrimination and enhanced separation of the four reporter populations compared with conventional cytometry (Figure 4A). This improved resolution enabled accurate FRET quantification for four previously unusable compounds: aminoresveratrol, salermide, AGK2, and BML266 (Figure 4B). In contrast, piceatannol, BML278, and sirtinol continued to interfere with fluorescence signals in the CT line and were therefore excluded. Among the validated compounds, the sirtuin inhibitors AGK2, salermide, and BML266 induced concentration-dependent increases in FRET efficiency in both TUB and VIM cells without affecting the CT control line (Figure 4C). Salermide additionally produced a modest increase in FRET in ACT cells (+8% at 100 µM). Spectral cytometry also enabled direct monitoring of intracellular compound accumulation based on intrinsic fluorescence intensity (Supplementary Figure 4C). Cells exhibiting the highest AGK2 fluorescence displayed enhanced FRET in TUB and VIM cells but not in CT cells (Figure 4D), indicating that intracellular drug accumulation correlated with cytoskeletal responses. Overall, spectral CytoFRET2 significantly improved barcode resolution and enabled reliable analysis of intrinsically fluorescent compounds. The validated sirtuin inhibitors produced effects like those observed with KDAC inhibitors, promoting stabilization of both microtubules and vimentin filaments.

**Figure 4.**
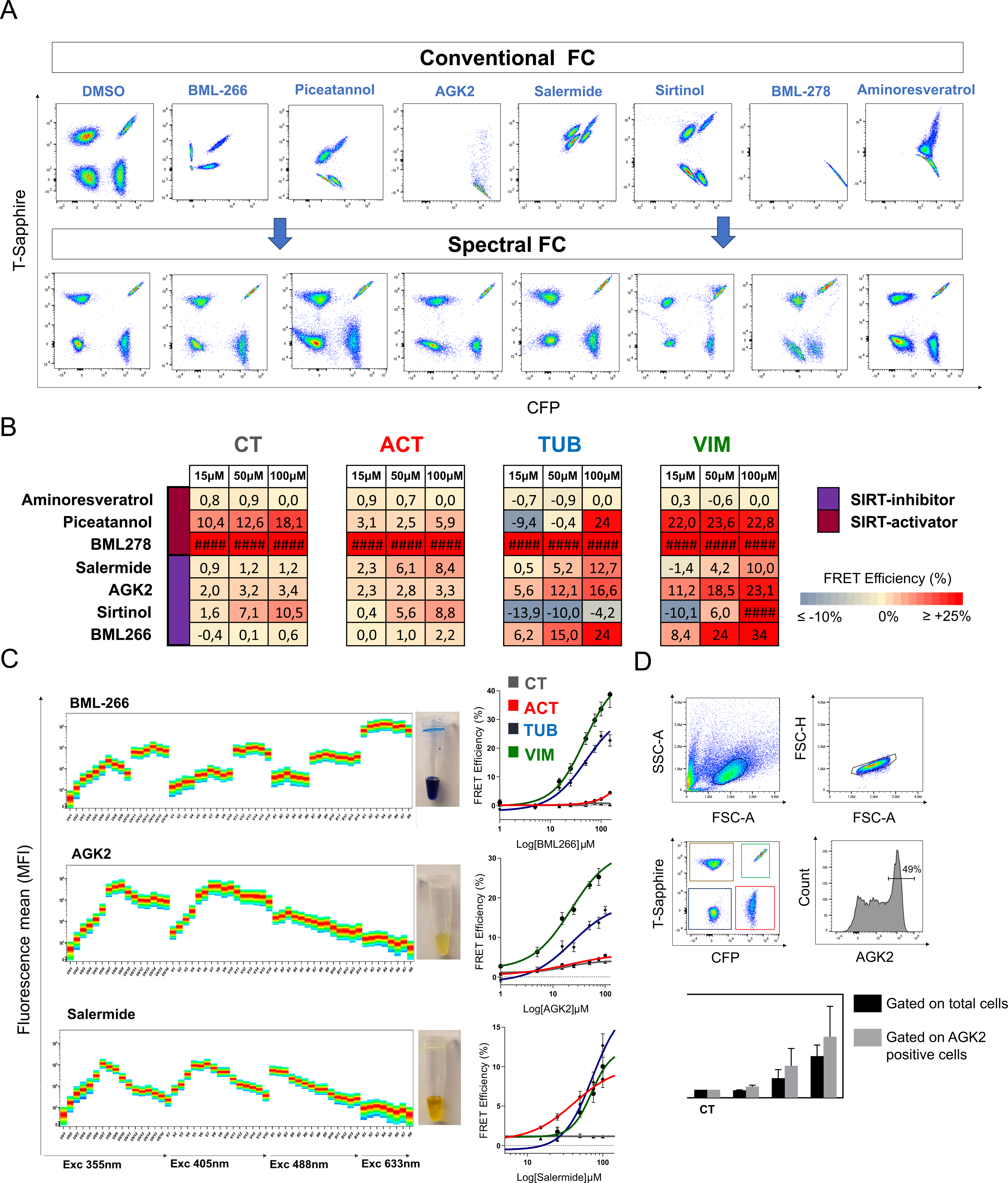
Spectral CytoFRET2 analysis of intrinsically fluorescent compounds. **(A)** Representative flow cytometry dot plots comparing fluorescent barcoding resolution obtained by conventional and spectral flow cytometry following treatment with intrinsically fluorescent compounds. **(B)** Heatmaps showing FRET efficiencies measured by spectral flow cytometry in ACT, TUB, VIM, and CT reporter lines following treatment with fluorescent compounds. Data represent three independent experiments performed in triplicate. **(C)** Spectral emission profiles of the fluorescent sirtuin inhibitors BML266, AGK2, and salermide, together with representative photographs of compound-containing tubes and corresponding FRET efficiency dose–response curves. **(D)** Correlation between AGK2 intracellular fluorescence intensity and FRET efficiency. Representative flow cytometry plots illustrate gating based on cell morphology and fluorescent barcoding. Histograms show selection of AGK2-high cells according to fluorescence intensity. Bar graphs compare FRET efficiencies between gated AGK2-high populations and total cell populations.

## DISCUSSION

In this study, we developed CytoFRET2, a multiplexed flow cytometry platform that enables real-time monitoring of cytoskeletal dynamics in living suspension cells. By combining genetically encoded FRET-based reporters with fluorescent cell barcoding, CytoFRET2 simultaneously interrogates the assembly status of the three major cytoskeletal systems—actin filaments, microtubules, and vimentin intermediate filaments—within a single mixed-cell population. This strategy substantially extends the capabilities of the original CytoFRET assay beyond actin dynamics ^22^ and provides a versatile framework for quantitative, high-throughput analysis of cytoskeletal organization in non-adherent cells. A major strength of CytoFRET2 lies in its compatibility with both conventional and spectral flow cytometry platforms. Although conventional flow cytometry allowed reliable detection of cytoskeletal remodeling events, spectral cytometry markedly enhanced assay performance by resolving fluorescence interference arising from intrinsically fluorescent compounds and by minimizing the impact of cellular autofluorescence associated with endogenous metabolic cofactors such as NADH and FAD ^31^. This advantage proved particularly valuable for the analysis of sirtuin inhibitors, a compound class frequently characterized by strong intrinsic fluorescence that compromises conventional flow cytometric measurements. Among the eight compounds that could not be reliably analyzed using conventional cytometry, approximately half became fully exploitable when spectral unmixing approaches were applied. The remaining compounds generated fluorescence signals sufficiently intense to saturate detector channels, thereby impairing accurate spectral deconvolution. Although these compounds could be analyzed at substantially lower concentrations, the concentrations required to avoid detector saturation were below those necessary to induce measurable biological effects. Similarly, reducing detector gains improved signal acquisition but simultaneously decreased the sensitivity required for optimal barcode discrimination and FRET signal resolution. These observations highlight a current technical limitation of spectral cytometry and suggest that future increases in detector dynamic range will further expand the applicability of multiplexed FRET-based screening approaches. An additional advantage of spectral analysis is the ability to directly monitor intracellular accumulation of fluorescent compounds, thereby providing a unique opportunity to correlate drug uptake with cytoskeletal responses at the single-cell level. Combined with the simultaneous acquisition of viability and morphological parameters, this feature considerably broadens the utility of CytoFRET2 for pharmacological screening and mechanistic studies.

Application of CytoFRET2 to a focused library of epigenetic modulators identified protein acetylation as a major regulatory mechanism governing cytoskeletal organization. Inhibition of KDACs, including both classical HDACs and sirtuins, promoted stabilization of microtubules and vimentin intermediate filaments, whereas inhibition of KATs induced the opposite phenotype on the vimentin network. These findings reveal coordinated regulation of multiple cytoskeletal systems by acetylation-dependent signaling pathways. The effects of HDAC inhibition on microtubules were largely anticipated. Numerous studies have demonstrated that inhibition of HDAC6 or SIRT2 increases α-tubulin acetylation, resulting in enhanced microtubule stability and resistance to depolymerization ^32,33^. Likewise, the almost complete absence of significant microtubule destabilization following KAT inhibition was expected. Indeed, the KAT inhibitors used in this study, including garcinol and anacardic acid, preferentially target the p300/CBP family of acetyltransferases ^34,35^, whereas α-tubulin acetylation is primarily catalyzed by αTAT1 ^36^. In contrast, the effects observed on the vimentin network were particularly intriguing. While HDAC inhibition promoted vimentin assembly, KAT inhibition induced a marked disassembly phenotype. Several proteomic studies have identified vimentin as an acetylated protein ^37^. However, the functional consequence of vimentin acetylation on filament assembly and network dynamics remains poorly understood. Our results provide direct evidence linking acetylation-dependent pathways to vimentin organization in living cells and suggest that reversible lysine acetylation may represent a previously underappreciated mechanism regulating intermediate filament dynamics.

Whether this regulation occurs through direct acetylation of vimentin or indirectly through other cytoskeletal components remains an important unresolved question. One plausible mechanism involves the extensive functional crosstalk between microtubules and vimentin filaments. Previous studies have shown that acetylated microtubules provide stable trafficking tracks for kinesin- and dynein-dependent transport of vimentin precursors and filament subunits ^12,38^. Furthermore, α-tubulin acetylation enhances motor protein processivity and intracellular transport efficiency ^39,40^. Increased microtubule acetylation could therefore promote homogeneous vimentin distribution and network stability, whereas reduced acetylation might impair vimentin trafficking, leading to filament fragmentation and disassembly. Such a model would provide a mechanistic framework linking acetylation-dependent regulation of microtubules to the observed remodeling of the vimentin cytoskeleton. Nevertheless, a direct role for vimentin acetylation cannot be excluded. Future studies combining quantitative mass spectrometry with site-directed mutagenesis will be required to identify acetylated lysine residues and determine their contribution to vimentin assembly. Likewise, genetic models deficient in α-tubulin acetylation, such as αTAT1-knockout cells or cells expressing non-acetylatable α-tubulin mutants, will be instrumental in determining whether the effects observed on vimentin are mediated primarily through microtubule acetylation or through direct modification of vimentin itself.

Collectively, these findings establish spectral CytoFRET2 as a robust and scalable platform for dissecting cytoskeletal regulatory networks in living cells and for high-content functional screening and mechanistic exploration of cytoskeletal biology. Beyond cancer biology, this approach should prove broadly applicable to studies of immune cell activation, mechanotransduction, stem cell biology, and pathological conditions characterized by extensive cytoskeletal remodeling. From a translational perspective, our results suggest that pharmacological modulation of acetylation may influence cellular plasticity through coordinated regulation of microtubules and vimentin, two cytoskeletal systems strongly associated with metastatic dissemination, epithelial-to-mesenchymal transition, and immune evasion. Finally, the previously unrecognized effects of KATs inhibitors on vimentin organization may open new opportunities for targeting aberrant cytoskeletal dynamics linked to pathological conditions, including cancer.

## Acknowledgments

We thank Frédéric Brau (IRCAN) and Julie Cazareth (IPMC) for their assistance with confocal imaging and spectral cytometry, respectively. The C3M imaging facilities for technical help. This work was supported by funds from Inserm, Ligue Contre le Cancer, Fondation ARC pour la recherche sur le cancer and Canceropôle Sud. The financial contribution of the Conseil général 06 and Région Sud to the C3M is also acknowledged.

## Author Contributions

F.L. and M.D. designed the study. F.L. performed the experiments and analyzed the data with the help of M.I. F.L. and M.D. wrote the original draft. M.D. supervised the study and edited the final version of the manuscript with the help of S.T.-D and F.L.

## Data Availability Statement

The raw data supporting the conclusions of this article are available by the authors upon request. Supplementary information accompanies this paper.

