## Supplemental files for "Spectral CytoFRET2 identifies lysine acetyltransferase inhibitors as modulators of vimentin assembly"

### Supplementary Table 1

| Name | CAS | Concentration | Compound description | Solvent |
|---|---|---|---|---|
| Trichostatin A | 58880-19-6 | 10mM | HDAC inhibitor | DMSO |
| 2,4-Pyridinedicarboxylic Acid | 499-80-9 | 10mM | Histone demethylase inhibitor | DMSO |
| Garcinol | 78824-30-3 | 10mM | HAT inhibitor | DMSO |
| Splitomicin | 5690-03-9 | 10mM | SIRT-2 inhibitor | DMSO |
| BML-210 | 537034-17-6 | 10mM | HDAC inhibitor | DMSO |
| Apicidin | 183506-66-3 | 10mM | HDAC inhibitor | DMSO |
| Suberoyl bis-hydroxamic acid | 38937-66-5 | 10mM | HDAC inhibitor | DMSO |
| Scriptaid | 287383-59-9 | 10mM | HDAC inhibitor | DMSO |
| Nullscript | 300816-11-9 | 10mM | Scriptaid Neg control | DMSO |
| 5-Aza-2'-deoxycytidine | 2353-33-5 | 10mM | DNA Me transferase inhibitor | DMSO |
| Zebularine | 3690-10-6 | 10mM | DNA Me transferase inhibitor | DMSO |
| SAHA | 149647-78-9 | 10mM | HDAC inhibitor | DMSO |
| Isonicotinamide | 1453-82-3 | 10mM | nicotinamide antagonist | DMSO |
| ITSA-1 | 200626-61-5 | 10mM | Inhibitor of TSA activity | DMSO |
| Phenylbutyrate·Na | 1716-12-7 | 10mM | HDAC inhibitor | DMSO |
| Tranylcypromine hemisulfate | 13492-01-8 | 10mM | Lysine demethylase inhibitor | DMSO |
| Valproic acid | 99-66-1 | 10mM | HDAC inhibitor | DMSO |
| EX-527 | 49843-98-3 | 10mM | SIRT1 inhibitor | DMSO |
| Resveratrol | 501-36-0 | 10mM | SIRT1 activator | DMSO |
| M-344 | 251456-60-7 | 10mM | HDAC inhibitor | DMSO |
| Nicotinamide | 98-92-0 | 10mM | SIRT inhibitor | DMSO |
| BML-266 | 96969-83-4 | 10mM | SIRT2 inhibitor | DMSO |
| Piceatannol | 10083-24-6 | 10mM | SIRT activator | DMSO |
| Fluoro-SAHA | 149648-08-8 | 10mM | HDAC inhibitor | DMSO |
| Valproic acid hydroxamate | 106132-78-9 | 10mM | HDAC inhibitor | DMSO |
| AGK2 | 304896-28-4 | 10mM | SIRT2 inhibitor | DMSO |
| Salermide | 1105698-15-4 | 10mM | SIRT inhibitor | DMSO |
| MC-1293 | 117378-93-5 | 10mM | HDAC inhibitor | DMSO |
| Anacardic acid | 16611-84-0 | 10mM | HAT inhibitor | DMSO |
| BIX-01294 | 935693-62-2 | 10mM | Histone methyl transferase inhibitor | DMSO |
| Butyrolactone 3 | 778649-18-6 | 10mM | HAT inhibitor | DMSO |
| CTPB | 586976-24-1 | 10mM | HATactivator | DMSO |
| Oxamflatin | 151720-43-3 | 10mM | HDAC inhibitor | DMSO |
| Sirtinol | 410536-97-9 | 10mM | SIRT inhibitor | DMSO |
| Suramin·6Na | 129-46-4 | 10mM | SIRT1 inhibitor | DMSO |
| BML-278 | 120533-76-8 | 10mM | SIRT1 actvator | DMSO |
| NCH-51 | 848354-66-5 | 10mM | HDAC inhibitor | DMSO |
| CI-994 | 112522-64-2 | 10mM | HDAC inhibitor | DMSO |
| Aminoresveratrol sulfate | 1224713-76-1 | 10mM | SIRT1 activator | DMSO |
| BML-281 | 1045792-66-2 | 10mM | HDAC-6 inhibitor | DMSO |
| Triacetylresveratrol | 42206-94-0 | 10mM | SIRT1 activator | DMSO |

**Supplementary Table 1.** Compounds included in the epigenetic library. The table lists the compound name, CAS number, solvent, stock concentration, and a brief description

**Supplementary Figure Legends**

**Supplementary Figure 1.** (A) Schematic representation of the lentiviral constructs used to generate reporter Jurkat T cell lines for monitoring actin, tubulin, and vimentin dynamics, together with the corresponding fluorescent barcoding strategy. The CT cell line expresses free CFP, T-Sapphire, GFP, and mOrange fluorescent proteins. The ACT cell line expresses GFP- and mOrange-tagged actin together with free CFP. The TUB cell line expresses GFP- and mOrange-tagged α-tubulin. The VIM cell line expresses GFP- and mOrange-tagged vimentin together with free T-Sapphire. (B) FRET efficiency dose–response curve following treatment with the actin-depolymerizing agent Latrunculin B. Mixed reporter cell lines were treated with increasing concentrations of Latrunculin B for 1 h at 37 °C.

**Supplementary Figure 2.** (A) Determination of cutoff values for the CT cell line following DMSO treatment. Twenty wells seeded with CT cells were treated in triplicate with DMSO at final concentrations of 15 µM, 50 µM, or 100 µM for 1 h at 37 °C. Heatmaps represent the percentage variation in FRET signal relative to the mean FRET signal obtained at the corresponding DMSO concentration. Cutoff values were defined based on the minimum and maximum observed variations. (B) FRET efficiency measured in reporter cell lines after 4 h treatment with 100 µM anacardic acid.

**Supplementary Figure 3.** Changes in the fluorescence emission spectra of GFP and mOrange associated with okadaic acid-induced FRET signals.

**Supplementary Figure 4.** (A) Fluorescence emission spectra of each fluorescent compound. Spectral flow cytometry histograms show the fluorescence emission profiles of Jurkat T cells treated with the indicated compounds. (B) Spectral flow similarity index between each fluorescent compound and the fluorescent proteins used in the assay. (C) Fluorescence intensity of each compound across all reporter cell lines. Flow cytometry histograms show the fluorescence intensity measured for each compound in all reporter cell lines.

### Supplementary Figure 1

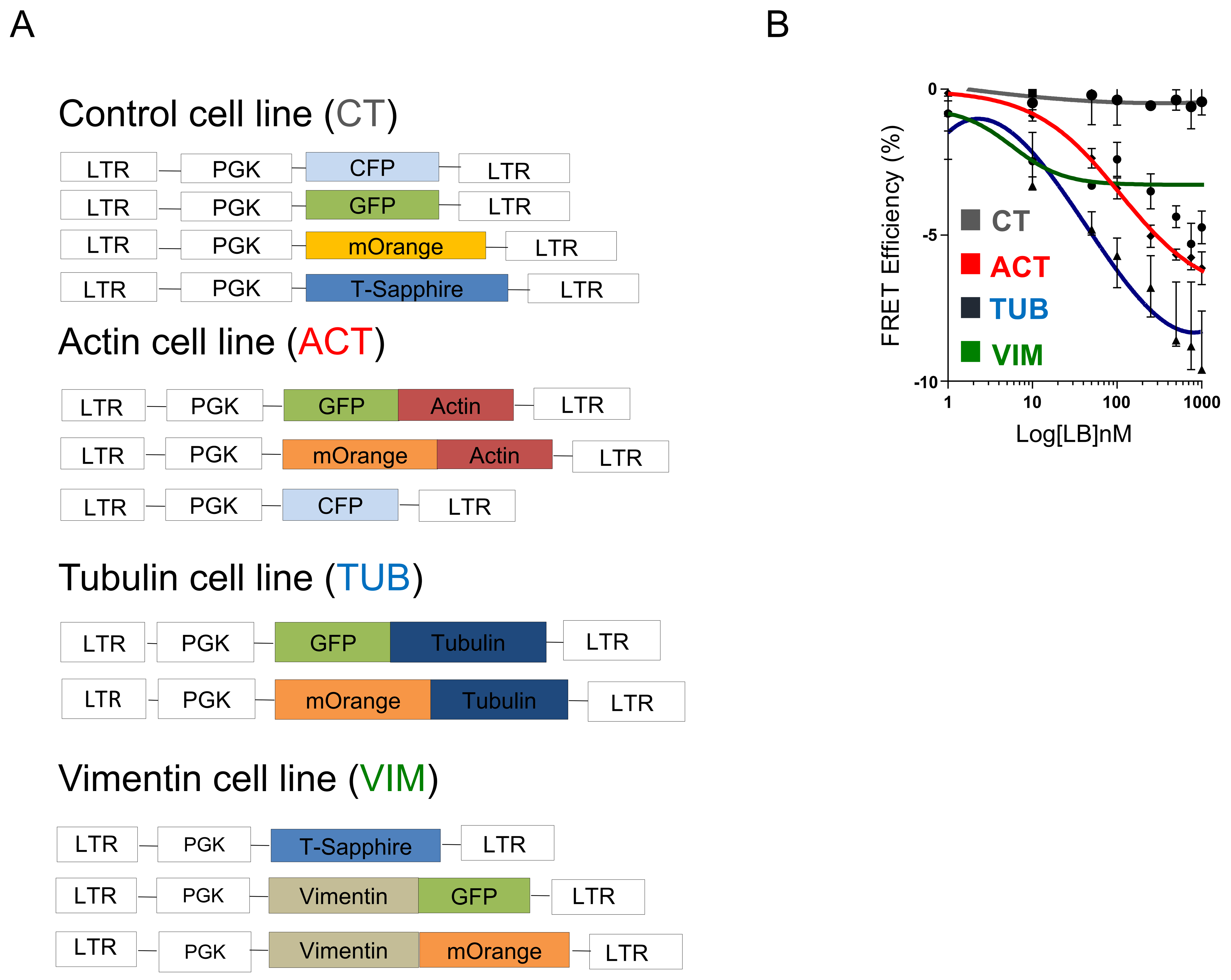

#### Supplementary Figure 2

A

| 100uM | | |
|---|---|---|
| 0,7 | 0,8 | 0,4 |
| 0,3 | 2,0 | 0,0 |
| 0,0 | 2,0 | 0,0 |
| 0,0 | 3,2 | 0,0 |
| 0,0 | -0,8 | -2,6 |
| -0,3 | -0,4 | 0,9 |
| 0,3 | 0,0 | 1,3 |
| 0,7 | 0,4 | -1,7 |
| 0,3 | -1,2 | 0,9 |
| 0,0 | -0,4 | 1,3 |
| 0,7 | 0,4 | 0,9 |
| 0,3 | -1,6 | 1,7 |
| -1,7 | 0,0 | -0,4 |
| -1,0 | 1,2 | 0,0 |
| -1,0 | 2,0 | -0,9 |
| -1,4 | 1,2 | -0,4 |
| -1,4 | -0,8 | -3,0 |
| -1,0 | -1,2 | -2,6 |
| -1,0 | -1,6 | -2,2 |
| -0,7 | -1,2 | -2,2 |
| **-3<cut off<3.2** | | |

| 50uM | | |
|---|---|---|
| -0,1 | 0,0 | 0,0 |
| -0,9 | -0,8 | 0,7 |
| -0,7 | -0,3 | 1,3 |
| -0,7 | 0,3 | 0,3 |
| -0,7 | -0,5 | -0,3 |
| -0,7 | 0,0 | 0,0 |
| -0,4 | 0,3 | 0,0 |
| -0,4 | 0,0 | 0,3 |
| -0,4 | 0,8 | 0,3 |
| -0,4 | 0,0 | 0,7 |
| 0,4 | 0,3 | 0,3 |
| 0,4 | 0,3 | 0,7 |
| 0,4 | -1,1 | -1,6 |
| 0,4 | -1,4 | -1,3 |
| 0,7 | -1,4 | -1,3 |
| 0,9 | -1,1 | -0,7 |
| 0,7 | -0,8 | -1,3 |
| 0,7 | 0,0 | -1,3 |
| 1,2 | 0,5 | -1,3 |
| 0,1 | 0,5 | -1,0 |
| **-1.6<cut off<1.3** | | |

| 15uM | | |
|---|---|---|
| -0,5 | -0,3 | 0,0 |
| 1,0 | 0,3 | 0,0 |
| 0,8 | 0,5 | 0,4 |
| 0,3 | 1,3 | -0,3 |
| 0,5 | 0,3 | 0,7 |
| 0,3 | 0,0 | 1,4 |
| 1,0 | 0,3 | 1,4 |
| 0,0 | 0,7 | 1,0 |
| 1,3 | -0,3 | 1,4 |
| 1,6 | 0,0 | 1,7 |
| 1,0 | 0,0 | 1,7 |
| -0,8 | 0,0 | 1,0 |
| -0,5 | -1,0 | -2,7 |
| -0,5 | -0,7 | -1,7 |
| -0,3 | -0,5 | -2,0 |
| -1,8 | -0,5 | -1,4 |
| -1,6 | -2,0 | -2,0 |
| -1,3 | -2,0 | -1,4 |
| -1,8 | -1,0 | -0,3 |
| 1,0 | 0,0 | -1,4 |
| **-2.7<cut off<1.7** | | |

B

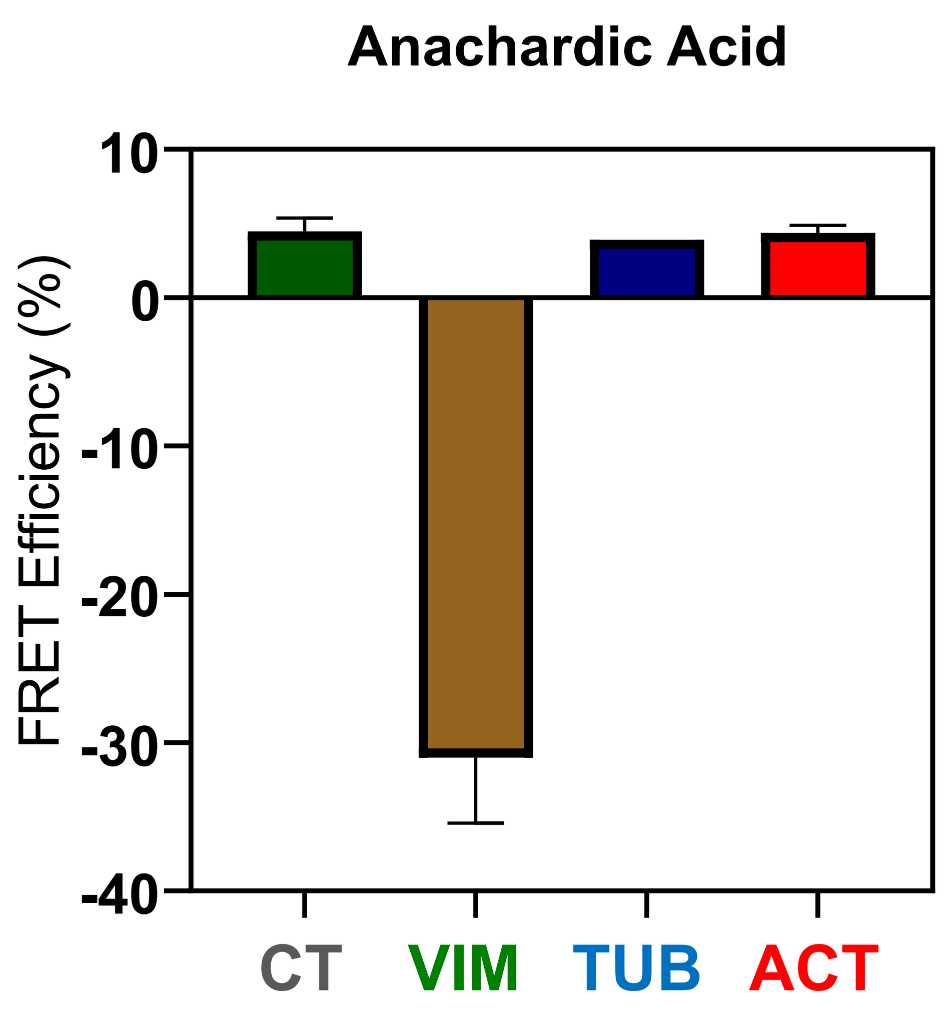

### Supplementary Figure 3

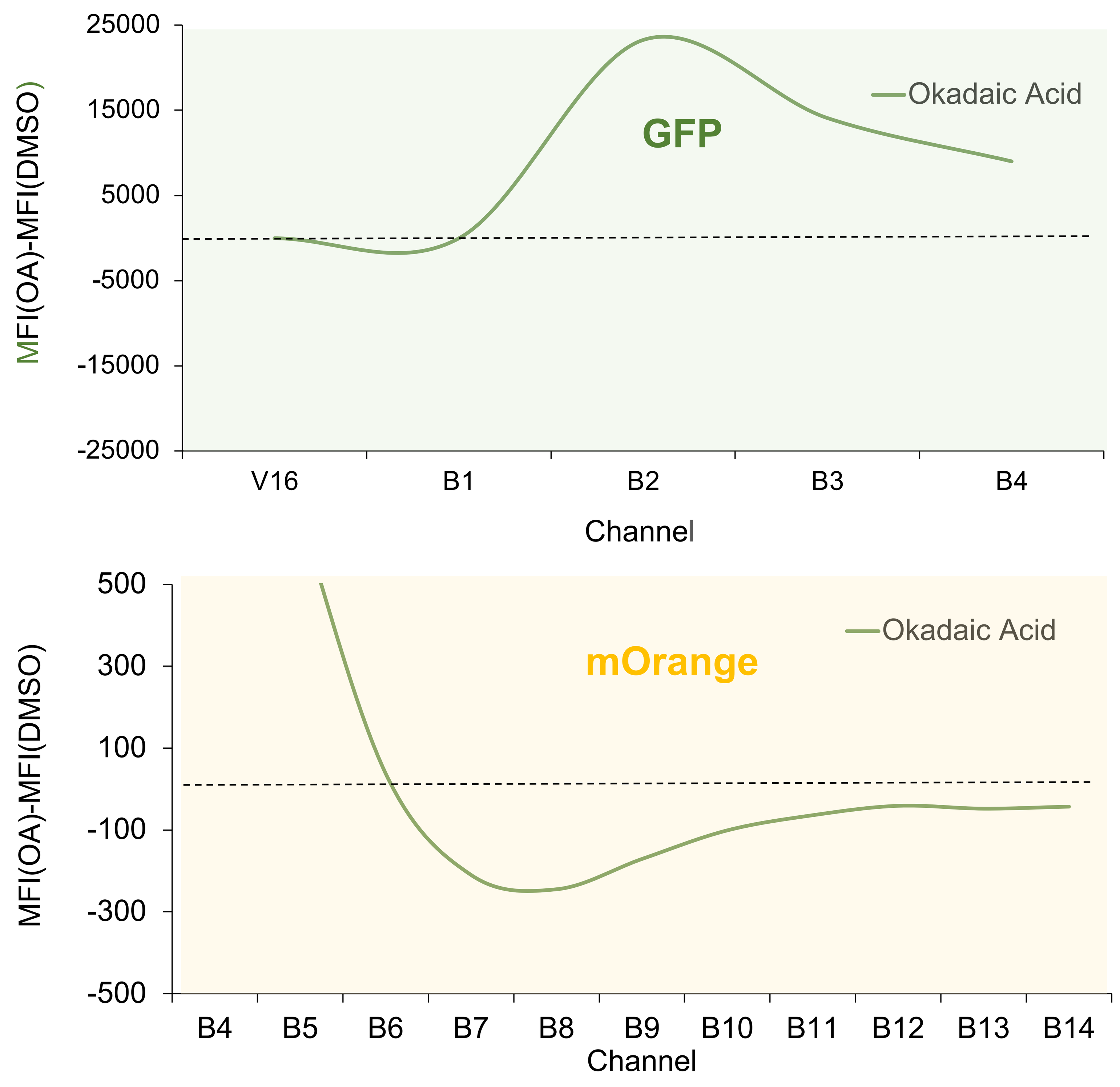

### Supplementary Figure 4

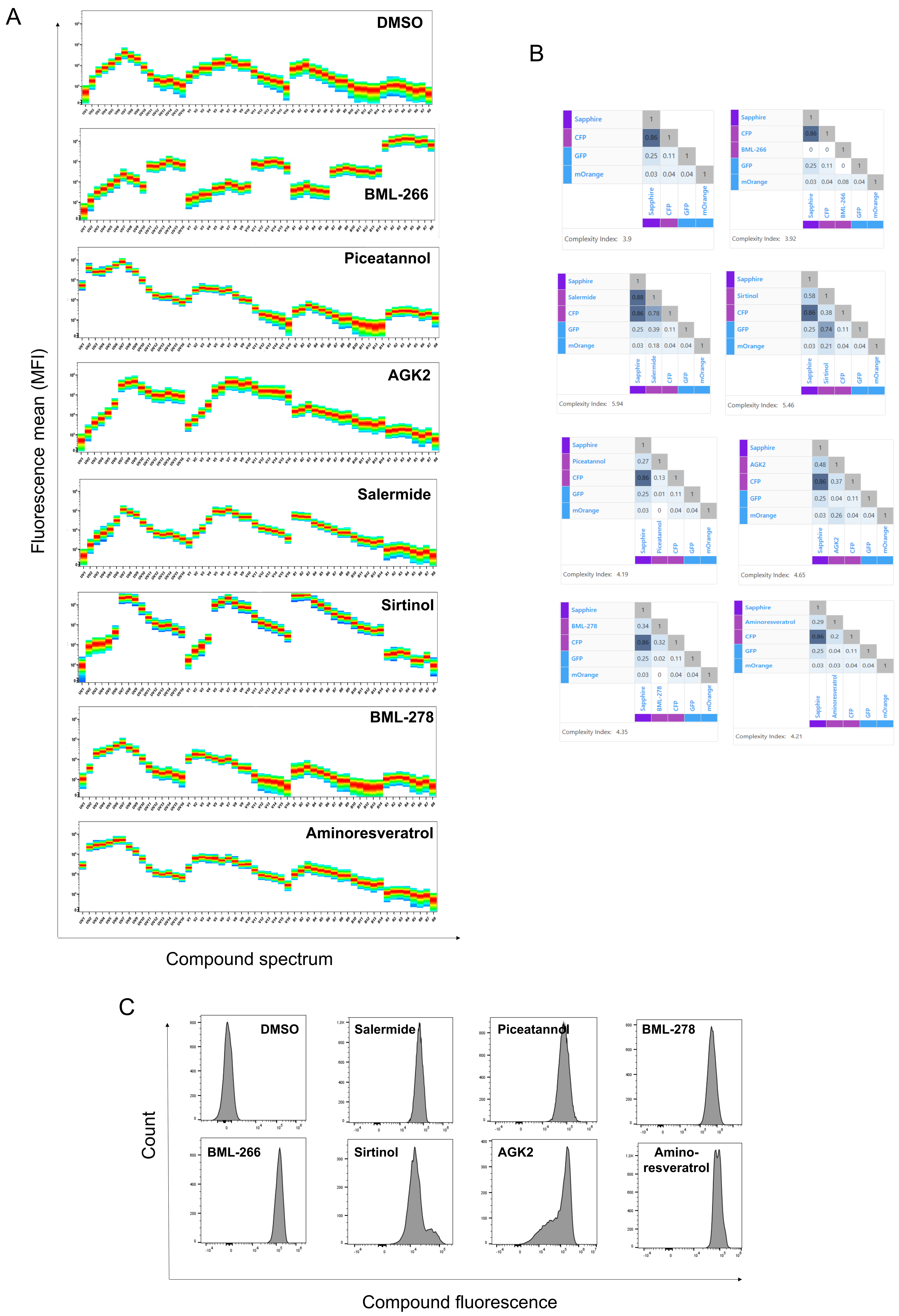
